# The extinction time under mutational meltdown

**DOI:** 10.1101/2022.02.01.478601

**Authors:** Lucy Lansch-Justen, Davide Cusseddu, Mark A. Schmitz, Claudia Bank

**Affiliations:** Instituto Gulbenkian de Ciência, Oeiras, Portugal; Institute of Evolutionary Biology, University of Edinburgh, United Kingdom; Grupo Física-Matemática, Faculdade de Ciências, Universidade de Lisboa, Portugal; Institute of Ecology and Evolution, University of Bern, Switzerland; Swiss Institute of Bioinformatics

**Keywords:** Mutational meltdown, lethal mutagenesis, extinction, mutagenic drugs, evolutionary theory

## Abstract

Mutational meltdown describes an eco-evolutionary process in which the accumulation of deleterious mutations causes a fitness decline that eventually leads to the extinction of a population. Possible applications of this concept include medical treatment of RNA virus infections based on mutagenic drugs that increase the mutation rate of the pathogen. To determine the usefulness and expected success of such an antiviral treatment, estimates of the expected time to mutational meltdown are necessary. Here, we compute the extinction time of a population under high mutation rates, using both analytical approaches and stochastic simulations. Extinction is the result of three consecutive processes: (1) initial accumulation of deleterious mutations due to the increased mutation pressure; (2) consecutive loss of the fittest haplotype due to Muller’s ratchet; (3) rapid population decline towards extinction. We find accurate analytical results for the mean extinction time, which show that the deleterious mutation rate has the strongest effect on the extinction time. We confirm that intermediate-sized deleterious selection coefficients minimize the extinction time. Finally, our simulations show that the variation in extinction time, given a set of parameters, is surprisingly small.

## 1 Introduction

The extinction of a population is a fundamental process in evolutionary biology and, given its irreversible nature, a lot of work across scientific fields has been devoted to its prediction (Ovaskainen and Meerson 2010; Carlson et al. 2014; Matuszewski et al. 2017; Wortel et al. 2021). For example, in medicine, the extinction time of a pathogen can decide whether its host survives or dies, whereas in conservation biology, dooming extinction calls for immediate action.

In asexual populations, one of the possible causes of extinction is related to excessively high mutation rates. Since the majority of mutations are deleterious, non-recombining populations can suffer from increasing mutation load when the mutation rate is high or the population size is low (or a combination of both). This process, which results in the step-wise successive loss of the group of individuals with the highest fitness in the population (the fittest class) due to the combined effect of mutation accumulation and genetic drift, is termed Muller’s ratchet (Muller 1964; Felsenstein 1974; Haigh 1978). In finite populations, Muller’s ratchet leads to a serial accumulation of deleterious mutations and, ultimately, extinction of the population. In evolutionary theory, this extinction process has been extensively studied under the names of mutational meltdown and lethal mutagenesis (Lynch et al. 1993; Bull et al. 2007; Elena and Sanjuán 2007; Matuszewski et al. 2017; Domingo and Perales 2019).

One promising application of the theory of mutational meltdown is the treatment of RNA virus infections (Bank et al. 2016; Ormond et al. 2017; Jensen and Lynch 2020; Jensen et al. 2020). That is because RNA viruses have exceptionally high mutation rates as compared to other viruses (Sanjuán et al. 2010), which may make them particularly susceptible to mutational meltdown by means of mutagenic drug treatment. Recently, mutagenic drugs such as Favipiravir and Molnupiravir have been developed that have shown promise for inducing mutational meltdown in various RNA viruses (Baranovich et al. 2013; Bank et al. 2016; de Avila et al. 2017). For instance, Favipiravir, a purine nucleoside analog, was proposed as a treatment for influenza viruses (Delang et al. 2018). Recent studies suggest that Molnupiravir, an analogue of the nucleoside cytidine, might be a promising tool against SARS-CoV-2 infections (Kabinger et al. 2021; Tao et al. 2021). One of the main concerns about the clinical use of mutagenic drugs is that, by enhancing mutation rates, the virus might be able to explore more genetic possibilities. These could include mutations that enable escape from mutational meltdown, or others that increase the success of the virus in future hosts the ones able to escape from the mutational meltdown (Nelson and Otto 2021). For instance, Bank et al. (2016); Goldhill et al. (2018) observed candidate mutations in influenza for resistance to Favipiravir in the lab. A crucial aspect in this context is the extinction (or survival) time of the virus under mutagenic treatment. Generally, the aim is that the mutagenic drug erases the virus population as quickly and predictably as possible. Therefore, it is fundamental to estimate the expected extinction time under the mutational meltdown process.

In this manuscript, we estimate the extinction time under mutational meltdown. We propose an analytical expression for the extinction time, which, to the best of our knowledge, has not been presented in the literature to date for the parameter regime considered here (described in section 2 Model and Methods). Using a simple model of a clonal non-recombining population allows for analytical calculations that are supported by simulations of the eco-evolutionary dynamics. Our estimates rely on the analysis of the three consecutive phases of the mutation accumulation and meltdown process, as described in Lynch et al. (1993): the rapid accumulation of mutations until the fittest class is lost, consecutive loss of the fittest class due to Muller’s ratchet, and the meltdown phase, in which the population rapidly goes to extinction. Our models predict how the mean time to extinction depends on the mutation rate, the wild-type reproduction rate, the deleterious fitness effect of mutations, and the carrying capacity of the population.

## 2 Model and Methods

We model the population dynamics and mutation accumulation of a non-recombining asexually reproducing population with a high mutation rate *μ*. The main variable of interest is the population size *N*, which varies over time. When *N* reaches zero, the population goes extinct. In our model the population size cannot exceed a given carrying capacity *K* imposed by the environment. We assume that all mutations have the same deleterious effect on fitness, represented by a selection coefficient –*s*, (*s* > 0). In particular, different mutations act independently on fitness and back mutations are neglected. As a measure for fitness we use the reproduction rate. The reproduction rate *w* as a function of the number of mutations *k* is given by

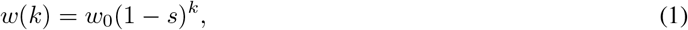

where *w*_0_ is the initial reproduction rate of the mutation-free population at time *t* = 0. Besides the population size, we also track the distribution of the number of mutations in the population, which we refer to as *mutation distribution* in the following. It determines the reproductive fitness of the population and indicates how the population size is changing. We model evolution in discrete time and with non-overlapping generations, starting with *N*_0_ founder individuals at time *t* = 0. Each generation, individuals obtain mutations and are replaced by their offspring. If necessary (i.e., if the new population size would exceed the carrying capacity), the population is sampled randomly until *K* individuals remain. Since mutations occur independently, the number of mutations an individual obtains each generation is approximately Poisson distributed with parameter *μ*. We assume that also the number of offspring of an individual with *k* mutations per generation is Poisson distributed with parameter *w*(*k*). Repeated cycles of mutation, reproduction, and population size control lead to an increasing mutation load and, hence, decreasing average population fitness. Since the carrying capacity ensures a finite population size, the unidirectional mutation accumulation leads to the eventual extinction of the population after the average fitness falls below one.

We are interested in biological systems in which extinction happens on short time scales, i.e. on the order of days to weeks as this time scale is most relevant for mutagenic drug treatments. Extinction happens on short time scales if the mutation-selection balance, as derived in Haigh (1978), is unstable, which results in the following condition on the model parameters

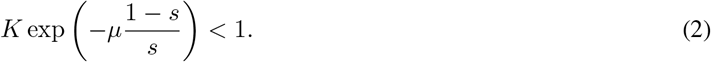

More details on the instability of the mutation-selection balance will be given in the next section 3.1.

We simulate the above-described population dynamics and mutation accumulation in an individual-based and stochastic model. In our simulations, we consider the following parameter regime: 1 < *w*_0_ ≤ 10,10^2^ ≤ *K* ≤ 10^4^, 10^−1^ ≤ *μ* ≤ 1 and 10^−3^ ≤ *s* ≤ 10^−1.5^. Smaller selection coefficients were not considered, as they yield very large extinction times. In the following, we present results from a mathematical analysis of this model, complemented by stochastic individual-based simulations. The complete annotated documentation of the computational analyses of this paper will be archived on Zenodo upon publication of the paper and is currently deposited at https://gitlab.com/evoldynamics/Time-to-mutational-meltdown.

## 3 Results

Our goal is to compute the time to extinction under the above-described model. It is determined by three successive phases of the evolutionary dynamics (Lynch et al. (1993), Figure 1). Initially, the population experiences rapid population size expansion and mutation accumulation (which we call the pre-ratchet phase). This is followed by a phase in which the fittest class of individuals is lost successively and at a constant rate while maintaining a constant population size (the ratchet phase). In the final phase (the meltdown phase), the population size collapses, resulting in the extinction of the population. In the following, we quantify the three phases separately, using different mathematical methods, and calculate the duration of each phase. The total time to extinction is given by the sum of the three time periods.

**Figure 1:**
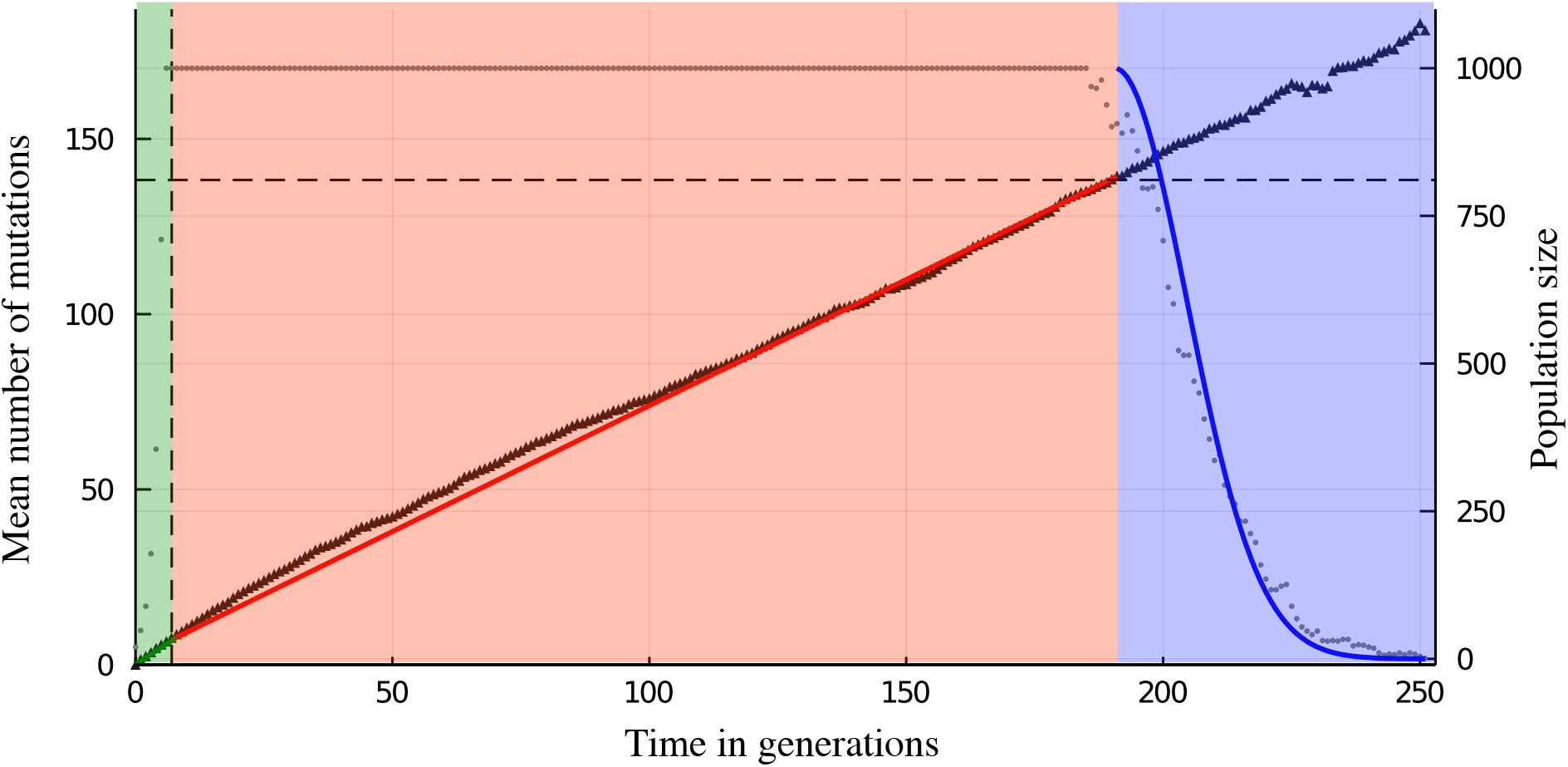
The mutational meltdown process consists of three phases, here indicated by regions shaded in different colors: initial rapid population growth and mutation accumulation towards the mutation-selection balance (left, green), followed by the ratchet phase with on average linear accumulation of mutations at the carrying capacity (middle, red), and, finally, the meltdown phase during which the population size rapidly decreases and mutations accumulate randomly (right, blue). Black dots represent the population size of one simulated population with founder population size *N*_0_ = 20, wild-type reproduction rate *w*_0_ = 2, mutation rate *μ* = 1.0, selection coefficient *s* = 0.005 and carrying capacity *K* = 1000. The black triangles represent the corresponding mean number of mutations in the population. The green line indicates the mean number of mutations according to equation (3). The red line shows the linear accumulation of mutations during the ratchet phase according to equation (6). The blue line indicates the decrease in population size according to equation (10).

### 3.1 Pre-ratchet phase

Starting from a monomorphic population without mutation load, the first phase of the evolutionary dynamics consists of rapid expansion of the population size and accumulation of deleterious mutations. We assume that the founder population size *N*_0_ is sufficiently large such that the population size reaches the carrying capacity quickly and the probability of both an early stochastic extinction and an early loss of the wild type can be neglected. If the population size is large, mean and variance of the number of mutations in the population are sufficient to describe the mutation distribution. Since both the number of newly accumulated mutations and the number of offspring per individual are Poisson distributed, the mutation distribution is also a Poisson distribution. At time *t* = 0 all individuals have zero mutations, which corresponds to the mutation distribution being ~ Poi(0). The mutation step shifts the distribution to the right by adding +*μ* to its parameter, whereas the reproduction step shifts it back to the left by multiplying with ·(1 – *s*); see Haigh (1978) for a detailed derivation. Population size control leaves the mutation distribution unchanged since individuals in excess are eliminated uniformly at random.

Successively repeating mutation and reproduction yields the mutation distribution as a function of time in generations

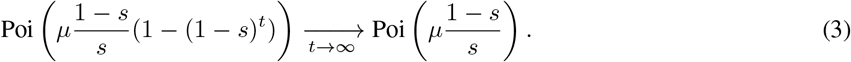

The mutation distribution approaches mutation-selection balance with parameter 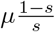. This deviates from the classical result derived in Haigh (1978) (mutation-selection balance with parameter 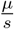) because we follow the distribution after reproduction instead of after mutation. The frequency of individuals with zero mutations is then

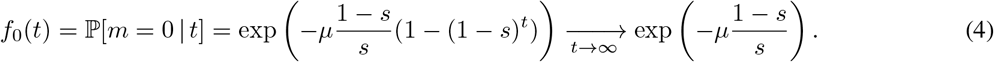

Consequently, the expected number of individuals with zero mutations in a population of size *K* is *n*_0_(*t*) = *K f*_0_(*t*). Due to condition (2) this implies that, as the mutation distribution approaches the mutation-selection balance, the zero-mutation class is likely to be lost, leading to the first click of the ratchet. This occurs at a time *T_R_*, when the size of the zero-mutation class drops below one:

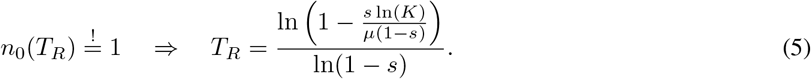

The mean number of mutations in the population at that time is 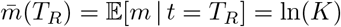.

### 3.2 Ratchet phase

Time *T_R_* heralds the ratchet phase that consists of repeated stochastic and successive loss of the fittest haplotype class. Consequently, the mean number of mutations in the population increases steadily. How fast the respective fittest class is lost on average, i.e. the speed of the ratchet, has been calculated in various ways and for different parameter regimes, see for example Haigh (1978); Pamilo et al. (1987); Lynch et al. (1993); Gabriel et al. (1993); Gessler (1995); Gordo and Charlesworth (2000b,a); Rouzine et al. (2008); Metzger and Eule (2013). The calculation of the ratchet speed is nontrivial as it depends on mutation, selection and genetic drift in an intertwined way. Moreover, the validity of the various derivations is strongly dependent on the parameter regime that is considered. We evaluated various proposed solutions and found that the expression derived by Gessler (1995) fitted our simulation data best, see Figure 2. However, Gessler’s formula exhibits a non-monotonic dependence on various parameters, which results from the discrete nature of its calculations. The speed of the ratchet depends on the selection coefficient and the difference between the number of mutations of the fittest class and the mean number of mutations in the population (the distance between the fittest class and the mean). Gessler calculated the fittest class and the mean of the mutation distribution (which Gessler determined to be a shifted negative binomial distribution) as discrete quantities, which depend on the model parameters *N, s* and *μ*. Varying the model parameters leads to discrete changes in fittest class and mean that occur at different parameter values for fittest class and mean respectively. Therefore, the distance between the fittest class and the mean exhibits a non-monotonic behaviour. Although the fittest class certainly is a discrete variable, the mean number of mutations in the population, in general, is continuous. The non-monotonicity of the ratchet speed is an artefact of this discrete approximation. In Appendix A, we derive a continuous extension of Gessler’s ratchet speed, (23) (see also Figure 2). Knowing the speed of the ratchet allows us to estimate the mean number of mutations in the population during the ratchet phase,

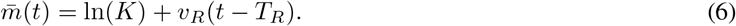

**Figure 2:**
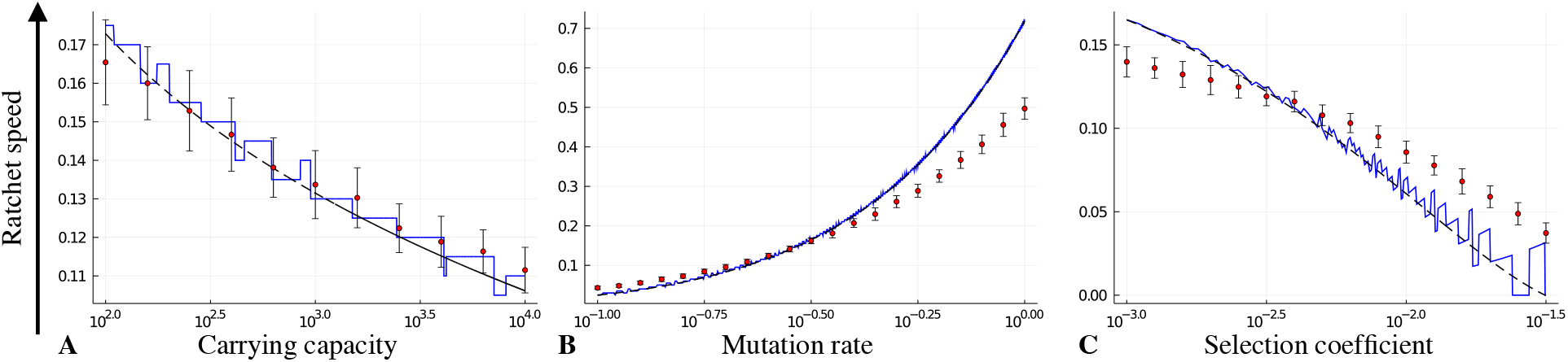
Estimation of the ratchet speed depending on the mutation rate, selection coefficient and carrying capacity. The ratchet speed decreases with increasing carrying capacity (A), increases with increasing mutation rate (B), and decreases with increasing selection coefficient (C). The expression for the ratchet speed derived in Gessler (1995) (blue solid line) is a good approximation. However, it is non-monotonic with respect to the relevant parameters. Our smooth approximation of Gessler’s ratchet speed (black dashed line) overcomes this limitation. Parameter values: founder population size *N*_0_ = 20, wild-type reproduction rate *w*_0_ = 2, selection coefficient *s* = 0.005, mutation rate *μ* = 0.27, carrying capacity *K* = 1000 and *n* = 50 simulation runs.

### 3.3 Meltdown phase

When the mean number of mutations in the population exceeds the critical threshold

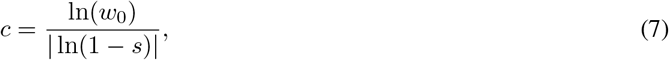

where *w*(*c*) = *w*_0_(1 – *s*)*^c^* = 1, the average fitness in the population drops below one. This is a turning point at which the meltdown phase begins. This happens at time *T_c_*, which we obtain from (6) as

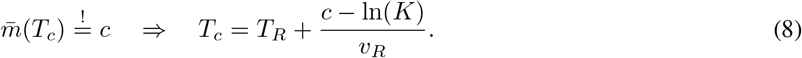

*T_c_* is given by the sum of two terms: the first, given in (5), is the time at which the ratchet starts, whereas the second represents the time until a population with a mean number of mutations 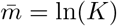 at *t* = *T_R_* reaches the critical value 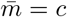, subject to a ratchet speed *v_R_*, given in (23).

During the meltdown phase, we assume selection to be inefficient. Hence, to get an estimate of the time length of this phase, we consider all individuals to obtain the same number of mutations and to have the same number of offspring each generation. In other words, we neglect genetic variation and describe the whole population by its mean number of mutations,

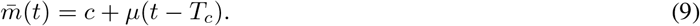

The population size as a function of time in generations can then be calculated recursively, *N*(*t* + 1) = *w*(*t*)*N*(*t*), starting at *N*(*T_c_*) = *K*, which yields

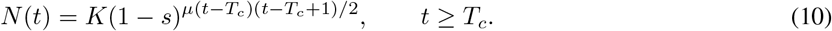

We assume that extinction occurs when the population size drops below one,

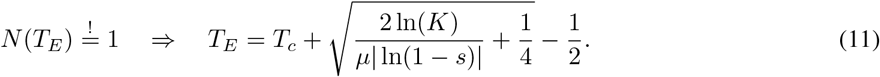

### 3.4 Extinction

Combining the three phases mentioned above yields the extinction time depending on the mutation rate *μ*, selection coefficient –*s*, wild-type reproduction rate *w*_0_ and carrying capacity *K*

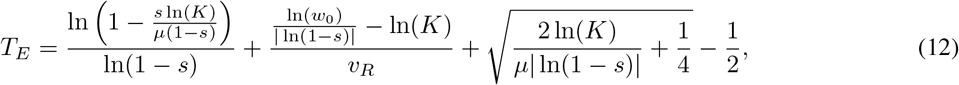

where the ratchet speed *v_R_* also depends on *μ*, –*s* and *K*. Note that the extinction time is independent of the initial population size *N*_0_, since we assume that the population size reaches the carrying capacity quickly.

The extinction time *T_E_*, given in (12), increases logarithmically with increasing wild-type reproduction rate *w*_0_, see Figure 3A. This is because a higher *w*_0_ increases the critical number of mutations (see equation (7)) which effectively shifts the process in time.

**Figure 3:**
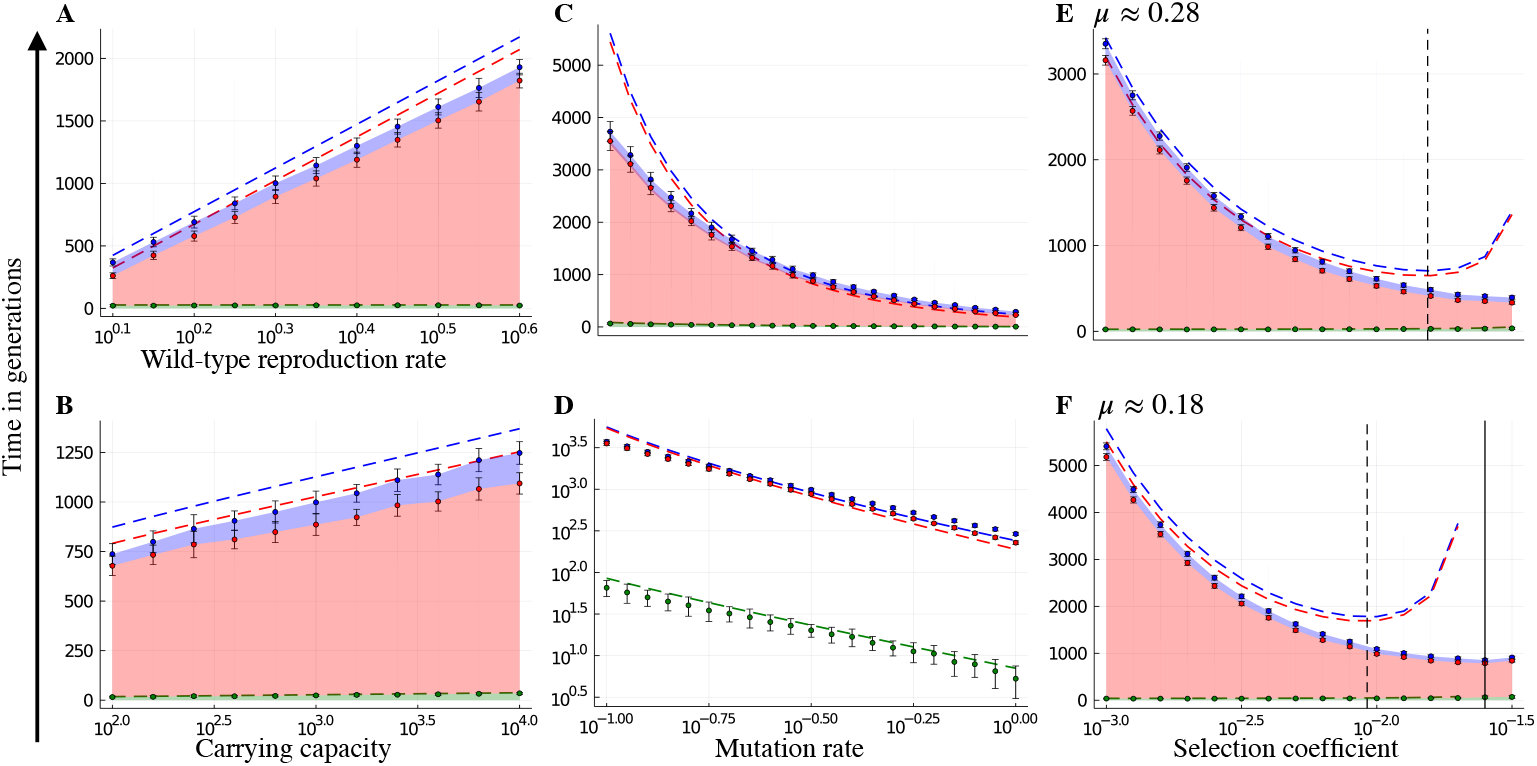
Duration of the three phases - pre-ratchet, ratchet and meltdown phase - depending on the wild-type reproduction rate, carrying capacity, mutation rate and selection coefficient. The phases are depicted by coloured ribbons that are stacked on top of each other. The pre-ratchet phase is given by the lowest ribbon in green, the ratchet phase in the middle in red and the meltdown phase on top in blue. Dots with error bars represent the mean *±* standard deviation obtained from stochastic simulations. The dashed lines are our analytical results, equations (5), (8) and (12). **A, B** The extinction time increases logarithmically with increasing wild-type reproduction rate and carrying capacity. Parameter values: *N*_0_ =20, *s* = 0.005, *μ* = 0.27, *K* = 1000 or *w*_0_ = 2 respectively and *n* = 50 simulation runs. **C, D** The extinction time decreases rapidly (according to a power law) with increasing mutation rate. Parameter values: founder population size *N*_0_ = 20, wild-type reproduction rate *w*_0_ = 2, selection coefficient *s* = 10^−2.3^ ≈ 0.005, carrying capacity *K* = 1000 and *n* = 50 simulation runs. **E, F** The extinction time is minimal for an intermediate selection coefficient. This optimal selection coefficient is higher for higher mutation rates and our predicted value (black dashed line) tends to underestimate the minimum value observed in simulations (black solid line). Parameter values: *N*_0_ = 20, *w*_0_ = 2, *K* = 1000, *μ* = 10^−0.55^ ≈ 0.28 or *μ* = 10^−0.75^ ≈ 0.18 respectively and *n* = 50 simulation runs.

Plotting the extinction time as a function of the carrying capacity *K* in a log-scale shows that *T_E_* increases logarithmically also with respect to *K*, see Figure 3B.

In contrast, the extinction time decreases rapidly with increasing mutation rate *μ*, see Figure 3C. This dependence arises because a higher mutation rate decreases the length of all three phases: it increases the speed towards the mutation-selection balance, the speed of the ratchet, and the speed of the meltdown process. Plotting the extinction time as a function of the mutation rate in a log-log scale results in an approximately linear decay see Figure 3D. This suggests that the extinction time decays as a power law of the mutation rate, i.e. *T_E_* ≈ *αμ*^-*β*^, for certain *α, β* > 0. We confirmed this relationship for different parameter sets, which showed variation in the exponent *β* as a function of the other three parameters. Unfortunately, our analysis of the relationship between extinction time and mutation rate as well as carrying capacity was limited to numerical solutions, because there exists no closed form of Gessler’s speed of the ratchet, *v_R_*, see equation (23) in Appendix A.

Consistent with previous results on Muller’s ratchet (Lynch et al. 1993), we found that the extinction time is minimal for an intermediate selection coefficient *s*\*, see Figure 3E, F and Figure 4A. A minimum at intermediate *s*\* is expected because extinction is a combination of different processes: When selection coefficients are small, many mutations can be accumulated without a large loss of fitness. When selection coefficients are large, selection is efficient at purging deleterious mutations. However, for intermediate selection coefficients, selection is less effective, yet every ratchet click leads to a significant fitness decrease. At higher mutation rates this minimum is attained at higher selection coefficients, compare Figure 3E, F.

**Figure 4:**
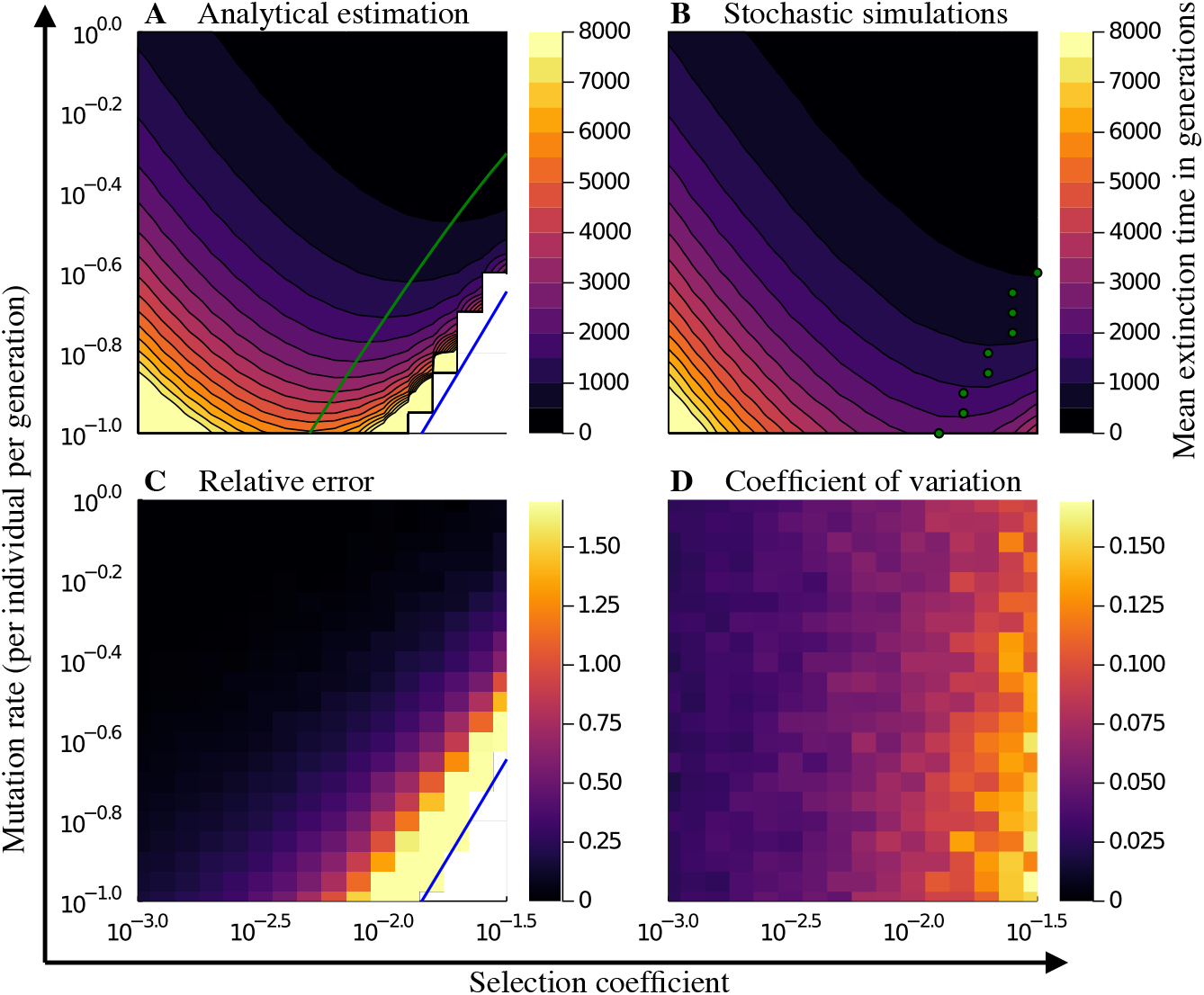
Comparing the analytically predicted mean extinction time with results from stochastic simulations for a range of mutation rates and selection coefficients. **A, B** The contour lines give parameter combinations with equal mean extinction time. Our analytical estimation is not applicable and returns an infinite extinction time if the mutationselection balance is stable, which is indicated by the blue line in panel A. This is not observed in stochastic simulations shown in panel B. The predicted optimal selection coefficient for which the extinction time is minimal (green line panel A) is smaller than the one observed in stochastic simulations (green dots, panel B). **C** The relative error of our analytical estimation is small in general but becomes large as the boundary of our parameter regime is approached. **D** The coefficient of variation of the extinction time in the simulations is comparably small and positively correlated with the selection coefficient. (Other parameters: founder population size *N*_0_ = 20, wild-type reproduction rate *w*_0_ = 2, carrying capacity *K* = 1000 and *n* = 50 simulation runs.)

### 3.5 Contributions of the three phases to the extinction time

Across the whole parameter range studied, the ratchet phase makes the largest contribution to the extinction time. When the parameters approach the boundaries of the parameter regime set by condition (2) (which happens for small mutation rates, large selection coefficients and large carrying capacities) the ratchet begins to click late, see equation (5), and the speed of the ratchet goes to zero. In contrast, the meltdown phase stays comparably short. Our simulation show that the ratchet phase remains the dominating phase also in this limit, see Figures 1 and 3.

In the case of small selection coefficients, the critical number of mutations becomes large, see equation (7), and so does the duration of the meltdown phase. Again, the ratchet phase remains the dominating phase, see Figure 3E, F.

### 3.6 Comparison with stochastic simulations

Comparing our analytical results with stochastic simulations shows that our expression is in good agreement with the extinction time of simulated populations at high mutation rates, see Figure 4. It accurately predicts the start of the ratchet, tends to overestimate the duration of the ratchet phase, and slightly underestimates the duration of the meltdown phase, see Figure 3. In total, our analytical expression, therefore, slightly overestimates the extinction time for most parameter combinations, see Figure 4C. In general, our estimate is the better the smaller the zero-mutation class under mutation-selection balance (left side of condition (2)). For example, for the parameters shown in Figure 3 (*N*_0_ = 20, *w*_0_ = 2, *s* = 10^−2.3^ ≈ 0.005 and *K* = 1000), the relative error ranges from ≈ 12.5% for intermediate mutation rates (*μ* = 10^−0.55^ ≈ 0.28) to ≈ 0.3% for high mutation rates (*μ* = 1.0). Note that our analytical approximation tends to predict a smaller optimal selection coefficient *s*\* than obtained from the simulations, see Figure 3E, F and Figure 4A, B. This is because the analytical expression is valid only in a certain parameter regime given by condition (2). At the boundaries of this regime the mutation-selection balance becomes stable, implying that the ratchet speed *v_R_* goes to zero and, therefore, the extinction time becomes infinite. For fixed carrying capacity and mutation rate, this happens when the selection coefficient becomes large (for example for *K* = 1000 and intermediate mutation rates, *μ* = 10^−0.55^ ≈ 0.28, at *s* ≈ 0.04 and for high mutation rates, *μ* = 1.0, at *s* ≈ 0.17). This explains the discrepancy between predicted and simulated optimal selection coefficient.

Interestingly, our simulations show that the variation in the extinction time is comparably small and correlates with the selection coefficient, see Figure 4D. For small selection coefficients, the variation is small as the mutation accumulation happens in many small steps, which averages out stochastic fluctuations. In contrast, for large selection coefficients, mutation accumulation is determined by rare and, hence, stochastic events, leading to a higher variation in the extinction times.

Figure 4 shows data with the carrying capacity of *K* = 1000, for data with *K* = 100 and *K* = 10000 see Figure S1 and S2.

## 4 Discussion

As a potential treatment option for SARS-CoV-2 infections, the use of mutagenic drugs against RNA virus infections has recently gained a lot of attention (Jensen and Lynch 2020; Jensen et al. 2020; Tao et al. 2021; Malone and Campbell 2021; Nelson and Otto 2021). The mode of action of such drugs is deeply rooted in evolutionary theory: most new mutations are deleterious and, therefore, an increase in the mutation rate can push a virus population to extinction, because selection is not efficient enough to weed out the deleterious variants. Importantly, unlike other drugs that attack virions individually, mutational meltdown is a population process that requires a strong and reliable action of the mutagenic drug. Specifically, extinction must occur as quickly as possible. That is because as the virus survives under the mutagenic pressure, it may not only accumulate deleterious mutations but also beneficial ones. Such mutations could help it survive in the presence of the drug (i.e., evolution of resistance), or they might be more general adaptations that make the virus more dangerous when it is transmitted to other hosts. Because of this danger that is specific to mutagenic drug treatments, it is important to theoretically know/predict and empirically minimize the expected time to population extinction under mutagenic treatment.

In this paper, we derive and analyze the mean extinction time of a population facing mutational meltdown under a simple model of population dynamics in a high mutation rate regime. As described in the Model and Results section, several simplifying assumptions underlie the mathematical analysis and the implementation of the stochastic simulations. We derive an analytical expression for the extinction time, equation (12), which allows us to determine how the extinction time depends on the model parameters: the mutation rate, the carrying capacity, the wild-type reproduction rate and the selection coefficient of mutations.

### 4.1 The mutation rate has the strongest effect on the extinction time

The extinction time decreases logarithmically with decreasing wild-type reproduction rate (Figure 3A) and with decreasing carrying capacity (Figure 3B). In contrast, the extinction time decreases much more rapidly (power-law dependence) with increasing mutation rate (Figure 3C, D).

The detected major effect of the mutation rate on the extinction time can be interpreted as a positive sign for the potential treatment of virus infections with mutagenic drugs. It indicates that the population dynamics and general initial fitness of the virus play a much weaker role than the mutation rate at determining whether and when the population will collapse under mutagenic drug treatment. This is important because we do not know the population dynamics inside the host and the reproductive rate of the virus when it enters the host, but we can possibly control the mutation rate of the virus by tuning the dosage of the mutagenic drug. Moreover, the power-law relationship between the mutation rate and the extinction time indicates that a small increase in dosage can result in a large decrease in the extinction time. Notably, this also applies in the reverse: if the dosage is only slightly too low, or if it does not sufficiently reach all body compartments in which the virus propagates, mutational meltdown may fail.

### 4.2 An intermediate selection coefficient minimizes the extinction time

Consistent with the previous literature (Lynch et al. 1993), we find that the extinction time is minimal for intermediate selection coefficients (Figure 3E, F). This minimum arises because larger selection coefficients lead to mutations with more severe negative effects on fitness, which speeds up the meltdown process. At the same time, larger selection coefficients make selection more efficient, which slows down the mutation accumulation. These two opposing effects result in an intermediate maximum of the rate of fitness decline under Muller’s ratchet (as found also for example in Gabriel et al. (1993)) that, in turn, minimizes the extinction time. This result points to an important limitation of our model: we assumed that the selection coefficient is constant, i.e., that every mutation has the same deleterious effect. Further work should evaluate whether the non-monotonicity of the extinction time with the selection coefficient holds when there is a distribution of selection coefficients.

The shown effectiveness of mutagenic drug treatments in experiments (Baranovich et al. 2013; Bank et al. 2016; Goldhill et al. 2019) suggests that the true distribution of selection coefficients of the virus is in a range that is indeed affected by Muller’s ratchet. This is consistent with experimental estimates of this distribution (Sanjuan et al. 2004; Jiang et al. 2016), which indicate that a large proportion of mutations have intermediately deleterious effects. Interestingly, the shifting minimum of the extinction time suggests that under different mutation rates, mutations with different selection coefficients could be the main contributors to the ratchet. It will be interesting to explore in future studies how the shift in the class of mutations that are most vulnerable to the ratchet can affect the observed mutation spectra during evolution in the presence and the absence of the drug, for example in laboratory evolution studies (Bank et al. 2016; Ormond et al. 2017).

### 4.3 The ratchet phase is the dominant contributor to the extinction time

Analyzing equation (12) allows us to determine how much the three different phases – pre-ratchet, ratchet and meltdown phase – contribute to the extinction time. We find that the ratchet phase is the dominating phase throughout the whole tested parameter range, see Figure 3. However, the duration of the ratchet phase is also the hardest to approximate because the ratchet speed depends on mutation, selection and genetic drift in a complex fashion. The ratchet speed is especially difficult to estimate in between regimes of a fast and slow-clicking ratchet, and no general theory that combines these regimes exists to date.

In the application of the theory to the case of mutagenic drug treatment, we are likely in a regime of a fast-clicking ratchet. Moreover, experimental measurements of the fitness effects of new mutations in this virus have shown that many mutations are of intermediate deleterious effect, which is important to keep the ratchet clicking (Sanjuan et al. 2004; Jiang et al. 2016). In this regime, Gessler’s approximation seems to be the currently best existing formula to compute the ratchet speed.

### 4.4 Stochastic simulations indicate small variation in extinction time

We performed stochastic simulations in order to test the validity of our analytical result and to quantify the variation in the extinction time under our model. We found that across a large range of the tested parameter regime, analytical expression and stochastic simulation are in good agreement, see Figure 4C. In our simulations, we observed mean extinction times ranging from around 65 generations for large mutation rates and large selection coefficients to 10000 generations for small mutation rates and small selection coefficients. The variation in the extinction time was surprisingly small, see Figure 4D. This suggests that the mean extinction time, which we derived in this paper, is a good predictor of the expected extinction time in simulations or experiments.

The observed small variation in extinction times is interesting to interpret in the context of application to mutagenic treatments. It suggests that when an experiment is repeated several times, or when many hosts are treated with the same dosage and under similar conditions, a prolonged extinction time may be an early, and easy-to-screen, signal of adaptation of the virus to the mutagenic drug treatment, and not just an expression of the stochasticity of the process. Future work should address in more detail how this observation holds in the presence of variable fitness effects and more complex demographic scenarios. However, we expect good robustness to these factors given that we found the ratchet phase to dominate the process and that the carrying capacity and the initial fitness contributed less to the extinction time than the mutation rate.

## Author contributions

LLJ and CB initiated the idea for the study. LLJ and DC derived the mathematical results. LLJ and MAS performed the simulations. LLJ, DC and CB drafted the paper. All authors critically revised the manuscript and gave approval of the final version for submission.

## Acknowledgements

The authors are grateful for helpful feedback on the manuscript received from Adamandia Kapopoulou and for discussion and inspiration provided by the THEE/Evoldynamics lab. This work was supported by ERC Starting Grant 804569 (FIT2GO), HFSP Young Investigator Grant RGY0081/2020 and EMBO Installation Grant 4152 to CB.

## Data Accessibility

The complete annotated documentation of the computational analyses of this paper will be archived on Zenodo upon publication of the paper and is currently deposited at https://gitlab.com/evoldynamics/Time-to-mutational-meltdown.

## Appendix A A smooth approximation of Gessler’s ratchet speed

Gessler (1995) approximated the rate of Muller’s ratchet (in particular, the rate of the loss of the fittest class) as

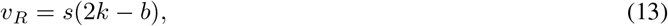

where

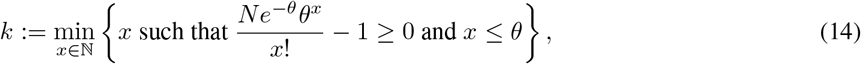

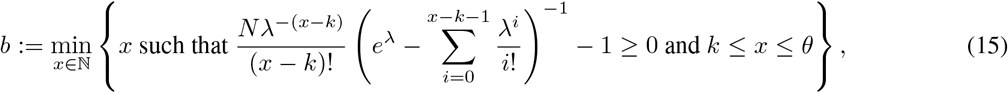

for *θ*:= *μ*(1 – *s*)/*s* and *λ*:= *θ* – *k*. Here N denotes the total population size, which Gessler assumed to be constant. In our model this translates to *N* = *K*, as our population remains at its carrying capacity during the ratchet phase.

For simplicity let us define

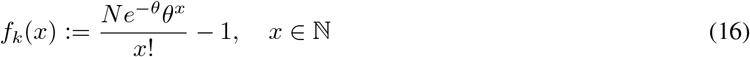

and

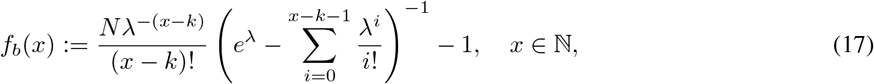

which appear in the definitions of the value *k* in (14) and *b* in (15).

Since *k*, 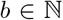, Gessler’s speed of the ratchet, eq. (13), can be non-monotonous with respect to the parameters, as shown in Figure 2. For instance, both coefficients *k* and *b* are monotonous and non-decreasing functions of the mutation rate *μ*. However, the following situation might occur:

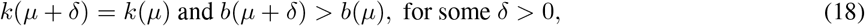

i.e., with increasing *μ*, *b* may take on a larger value while *k* remains constant, which results in a sudden decrease of the ratchet speed (which should monotonously increase with the mutation rate),

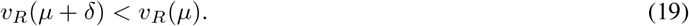

To avoid this behavior, we extend the domain of the functions *f_k_, f_b_* as defined in (16)-(17), to positive real numbers. We define

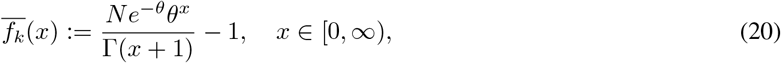

where we use the Euler Gamma function Γ which generalises the factorial operation over the real numbers (Abramowitz and Stegun 1972). In particular, 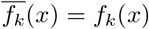 for all 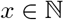.

The extension of the function *f_b_*(*x*) in (17) to positive real numbers needs more care because *x* appears in the upper bound of summation of 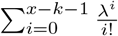. Defining 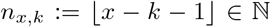, i.e. *n_x, k_* ≤ *x* – *k* – 1 < *n_x, k_* + 1, the function

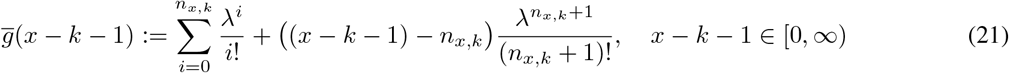

represents an interpolation of 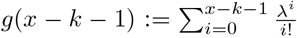 on [0, ∞). Indeed, when 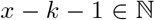, equation (21) corresponds to the original sum, and 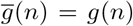 for all 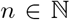. Therefore, the following function extends *f_b_* to positive real numbers:

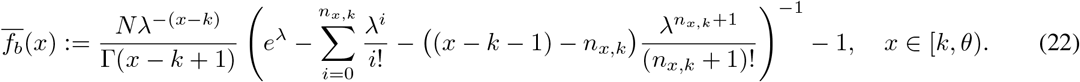

Since both *f_k_* and *f_b_* are increasing functions of 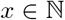, the problem of finding the minimum integers *k, b* such that *f_k_* (*k*), *f_b_*(*b*) ≥ 0, as required in equations (14) and (15), easily extends to the problem of finding the zeros of 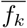 and 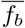. Therefore, if there exist 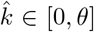 and 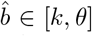, such that 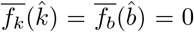, the following expression provides a continuous extension of the original Gessler’s ratchet speed:

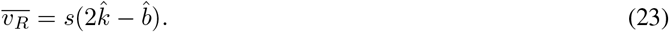

Similar to the original formula by Gessler (1995), the above expression is well defined only for suitable choices of the parameters *μ, s, N*.

Figure 2 in the main text compares the ratchet speed in its original discrete version by Gessler (1995) with our smooth monotonic extension.

