## Supplemental Data for "The extinction time under mutational meltdown"

### SUPPLIMENTARY MATERIAL

#### 1. Data for carrying capacities $K = 100$ and $K = 10000$

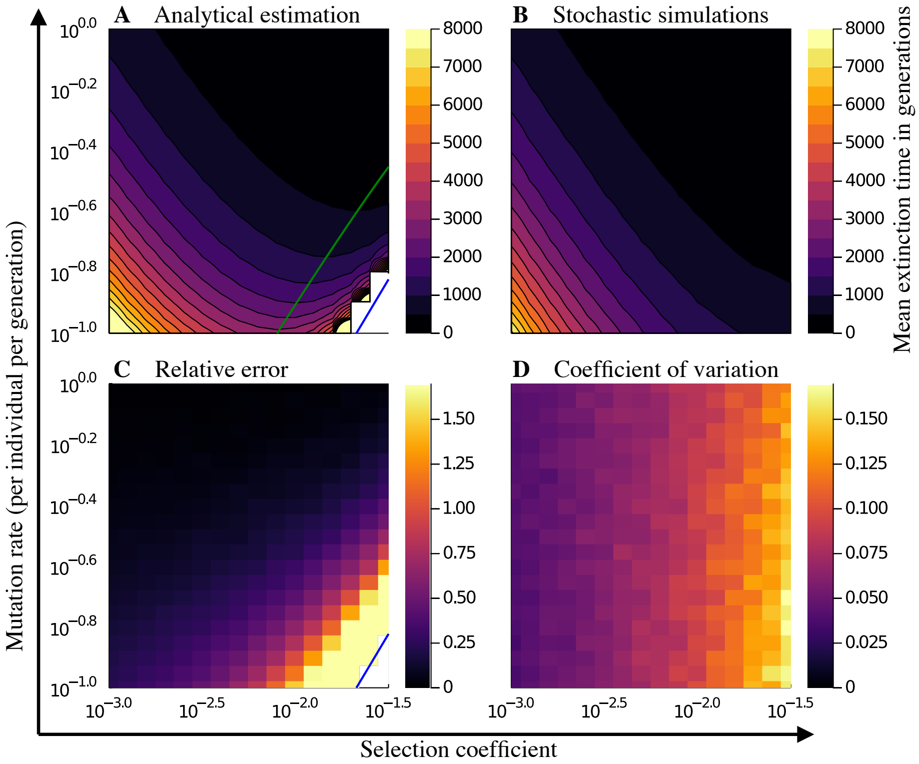


FIGURE 1. Carrying capacity $K = 100$: *Comparing the analytically predicted mean extinction time with results from stochastic simulations for a range of mutation rates and selection coefficients.* **A, B** The contour lines give parameter combinations with equal mean extinction time. Our analytical estimation is not applicable and returns an infinite extinction time if the mutation-selection balance is stable which is indicated by the blue line in panel A. This is not observed in stochastic simulations shown in panel B. The predicted optimal selection coefficient for which the extinction time is minimal (green line panel A) is smaller than the one observed in stochastic simulations. **C** The relative error of our analytical estimation is small in general but becomes large as the boundary of our parameter regime is approached. **D** The coefficient of variation of the extinction time in the simulations is comparably small, but positively correlated with the selection coefficient. (Other parameters: founder population size $N_0 = 20$, wild-type reproduction rate $w_0 = 2$ and $n = 150$ simulation runs.)

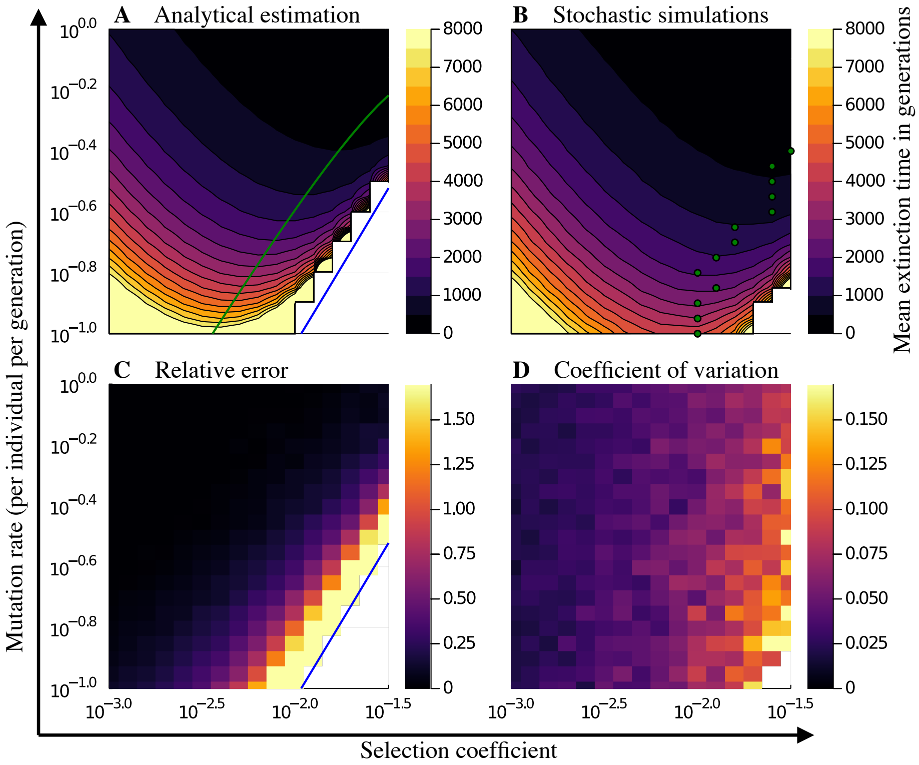


FIGURE 2. Carrying capacity $K = 10000$: *Comparing the analytically predicted mean extinction time with results from stochastic simulations for a range of mutation rates and selection coefficients.* **A, B** The contour lines give parameter combinations with equal mean extinction time. Our analytical estimation is not applicable and returns an infinite extinction time if the mutation-selection balance is stable which is indicated by the blue line in panel A. This is not observed in stochastic simulations shown in panel B. The predicted optimal selection coefficient for which the extinction time is minimal (green line panel A) is smaller than the one observed in stochastic simulations (green dots panel B). **C** The relative error of our analytical estimation is small in general but becomes large as the boundary of our parameter regime is approached. **D** The coefficient of variation of the extinction time in the simulations is comparably small, but positively correlated with the selection coefficient. (Other parameters: founder population size $N_0 = 20$, wild-type reproduction rate $w_0 = 2$ and $n = 20$ simulation runs.)
